# Stimulus Sampling Shapes Estimated Perceptual Geometry in Psychophysical Scaling

**DOI:** 10.64898/2026.10.30.763047

**Authors:** Clement Abbatecola, Kim Beneyton, Kenneth Knoblauch

**Author notes:** equal contributing authors (Clement Abbatecola, Kim Beneyton).

## Abstract

We used simulations and mathematical analysis to investigate how simultaneous estimation of transducer nonlinearity and perceptual geometry affects bias in maximum likelihood difference scaling (MLDS). Uniform sampling in the stimulus domain disproportionately sampled slowly varying regions and inadequately sampled rapidly varying regions of nonlinear transducers, leading to biased estimates of transducer shape. These biases were substantially reduced when stimuli were sampled uniformly in response space, distributing observations more evenly across the transducer. Likelihood-surface analysis revealed substantial confounding between sampling-induced distortions and effects attributable to changes in perceptual geometry. Uniform stimulus-domain sampling produced a pronounced trade-off between the parameters governing transducer nonlinearity and perceptual geometry, which was substantially reduced under uniform response-space sampling. Although response-space sampling generally improved the local conditioning of the likelihood surface, parameter identifiability remained dependent on its intrinsic geometry and thus on the underlying transducer. These results show that stimulus sampling can influence whether departures from a Riemannian perceptual metric are detected, highlighting the importance of sampling designs that adequately represent perceptual space. More generally, they demonstrate how appropriate sampling can help disentangle the effects of nonlinear transduction from those of nonlinear contributions of perceptual differences to judgments.

## Introduction

A central goal of psychophysics is to characterize how changes in physical stimuli are transformed into changes in perceived appearance. This relationship is commonly described by a psychophysical transducer function that maps physical stimulus values onto an internal perceptual representation [Stevens, 1957].

Classical threshold methods estimate the local slope of the transducer function through just-noticeable differences, relying on a local linear approximation [Knoblauch, 1993, May and Solomon, 2013]. Alternative approaches, including matching and magnitude estimation, seek to recover perceptual scales more directly but rely on assumptions that have been debated extensively [Marks, 1974]. Matching methods raise questions concerning the uniqueness of the resulting scales [Vincent et al., 2024], whereas magnitude estimation depends on assumptions about the relationship between numerical judgments and perceived magnitude [Devinck and Knoblauch, 2023, Zhou et al., 2024].

Maximum Likelihood Difference Scaling (MLDS) and Maximum Likelihood Conjoint Measurement (MLCM) instead estimate latent perceptual scales from ordinal comparisons of stimulus intervals rather than from thresholds or direct magnitude judgments [Maloney and Yang, 2003, Ho et al., 2008, Knoblauch and Maloney, 2008, Kingdom and Prins, 2016, Maloney and Knoblauch, 2020, Vacher and Mamassian, 2024]. Because these methods infer the perceptual representation indirectly from comparisons among stimuli, the choice of stimulus sampling strategy can influence the representation that is recovered and, potentially, the apparent geometry of perceptual differences.

MLDS assumes a set of *q* physical stimuli, (*ϕ*_1_, *ϕ*_2_, …, *ϕ*_*q*_), ordered along a physical continuum. The psychophysical task can be performed with 3 (triads) or 4 (quads) stimuli on each trial. To simplify, the triad method will be described. On each trial, three stimuli from the set are selected, (*ϕ*_*a*_, *ϕ*_*b*_, *ϕ*_*c*_), with *ϕ*_*a*_ *< ϕ*_*b*_ *< ϕ*_*c*_. The observer judges whether *ϕ*_*a*_ or *ϕ*_*c*_ is perceptually closer to *ϕ*_*b*_, or equivalently whether the interval between *ϕ*_*a*_ and *ϕ*_*b*_ is smaller than that between *ϕ*_*b*_ and *ϕ*_*c*_. If we assume that each *ϕ*_*i*_ evokes a perceptual response, *ψ*_*i*_ then the decision variable can be expressed as

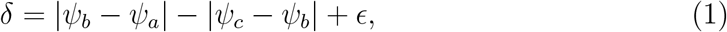

where *ϵ* is independent Gaussian judgment noise assumed to have mean 0 and variance *σ*^2^ [Maloney and Yang, 2003]. The judgment noise, *ϵ*, accounts for variability in responses when the two intervals are nearly equal. The subject chooses *ϕ*_*a*_ when *δ <* 0 and *ϕ*_*c*_ otherwise. This is an equal-variance Gaussian signal detection model [Green and Swets, 1966, Maloney and Yang, 2003]. Equation 1 will be referred to hereafter as the *standard model*. When the stimuli are ordered on the physical scale as above, the terms of the decision variable can be combined to yield,

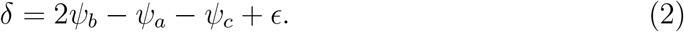

The scale values that best predict the observer’s responses are estimated by maximum likelihood, and since the decision variable is linear in the estimated coefficients, the fitting procedure can be implemented as a Generalized Linear Model (GLM) with a binomial family and probit link function [Knoblauch and Maloney, 2008]. The estimated scale has interval properties: equal changes in scale represent equal perceptual differences, but only its relative shape is uniquely determined, with translation and positive linear scaling remaining arbitrary [Krantz et al., 1971, Maloney and Yang, 2003, Knoblauch and Maloney, 2008].

In many applications of MLDS, the stimulus values are selected to be equally spaced in the physical stimulus domain. However, when the physical-to-perceptual transformation is nonlinear, equal spacing in the stimulus domain does not correspond to equal spacing in perceptual space. Within a family of power transducer functions, the estimated scale has been shown to be unbiased over powers spanning the range 0.25 to 4 [Maloney and Yang, 2003, Aguilar et al., 2026]. In addition, the shape of the transducer function estimated by MLDS was found to be robust to several forms of model misspecification, including violations of assumptions about the distribution and scaling of internal noise [Maloney and Yang, 2003, Kingdom, 2016, Aguilar et al., 2026]. These findings establish the robustness of MLDS to several departures from its modeling assumptions, but they do not address how the sampling of a nonlinear transducer affects estimation when transducer nonlinearity and perceptual geometry are estimated simultaneously.

Bujack and colleagues [Bujack et al., 2022, Teti et al., 2022] proposed a generalized decision variable in which observers may compare perceptual intervals through a nonlinear transformation rather than directly comparing their magnitudes. Specifically, they suggested replacing the partial perceptual differences in the standard MLDS decision rule with a compressive power transformation with exponent *α*, such that

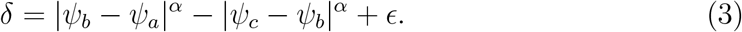

When 0 *< α <* 1, larger perceptual intervals contribute proportionally less to the decision variable, producing what the authors termed “diminishing returns”. They interpreted this departure from linear interval comparison as evidence that perceptual distances may not obey the usual assumptions of Riemannian geometry. However, the evidence for including diminishing returns in the decision model has been questioned on methodological grounds [Brainard, 2022].

Under the standard MLDS decision model, previous simulation studies have shown that MLDS can recover transducer shapes with little bias over a broad range of power nonlinearities [Maloney and Yang, 2003, Aguilar et al., 2026]. However, these results assume that the stimulus sampling strategy provides adequate coverage of the perceptual scale. If systematic biases arise from interactions between the physical-to-perceptual transducer and the experimental design, apparent deviations from the standard model could be inferred even when the underlying comparison process is linear. This possibility is particularly important when testing extensions such as the diminishing-returns model, in which departures from the standard model are interpreted as evidence for nonlinear interval comparison. Thus, apparent non-Riemannian geometry could in principle reflect inadequate sampling of a nonlinear transformation between the physical stimulus and the representation being compared, rather than a property of the underlying comparison process.

In this paper, we use Monte Carlo simulations to investigate whether stimulus sampling can produce apparent departures from perceptual geometry in MLDS. We examine the joint estimation of transducer nonlinearity and the diminishing-returns exponent across different transducers, perceptual geometries, and stimulus-sampling strategies. We show that uniform sampling in the physical stimulus domain introduces systematic bias and confounds the two parameter estimates, whereas sampling approximately uniformly in perceptual space substantially reduces both effects. More generally, the results demonstrate that apparent deviations from a proposed perceptual geometry can arise when a nonlinear perceptual transformation is inadequately sampled, emphasizing that experimental design is part of the inferential problem when perceptual scales and their geometry are estimated from ordinal comparisons.

## Methods

### Simulation design

All simulations were performed using the open source software R [R Core Team, 2026].

#### Power functions

The principal transducer functions investigated were drawn from a family of power functions,

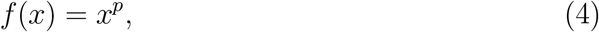

with 7 values of the exponent *p* ranging from 0.125 to 8 and spaced by factors of 2. These values span a range of compressive (*p <* 1) and expansive (*p >* 1) nonlin-earities. For each value of *p*, 4 values of the diminishing-returns exponent *α* were tested, ranging from 0.125 to 1 and spaced by factors of 2. For all simulations, the decision variable was calculated using the generalized difference rule in Equation 3, where *α <* 1 produces compression of larger perceptual differences. The case *α* = 1 corresponds to the standard model (Equation 1) with no diminishing-returns transformation.

The stimulus levels consisted of 11 values sampled from the interval (0, 1), either uniformly spaced in stimulus units (USS, Figure 1a) or uniformly spaced along the response scale (URS, Figure 1b). In each simulation, a session consisted of 5 repetitions of the 11!*/*(3! 8!) = 165 unique triads formed from the 11 stimulus levels, for a total of 825 trials.

**Figure 1.**
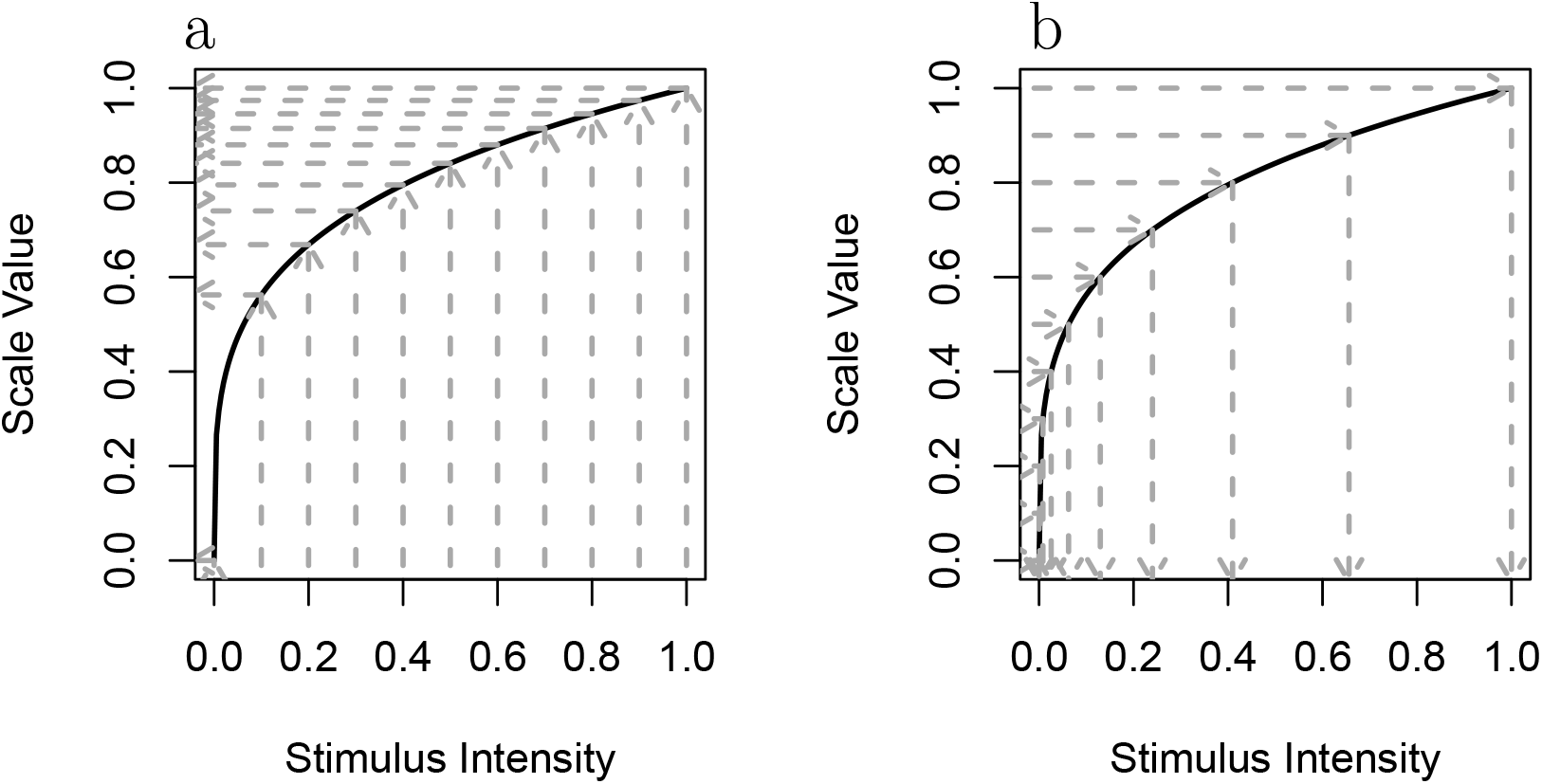
Uniform sampling in stimulus and response spaces. The solid curve in both graphs displays an example transducer function that is nonlinear in stimulus intensity. (a) The gray arrows map equally spaced stimulus intensities to unequally spaced responses. (b) The gray arrows map equally spaced responses to unequally spaced stimulus intensities.

For each trial, the decision variable, *δ*, from Equation 3 was computed by applying the transducer function to the three stimulus values in that trial. Gaussian random noise with mean 0 and standard deviation 0.15 was then added to *δ*, a noise level consistent with values typically observed in experimental data [Knoblauch and Maloney, 2012, Devinck and Knoblauch, 2012]. The choice was assigned a value of 0 or 1 according to whether the resulting decision variable was negative or positive, respectively. Each replication used an independently generated set of Gaussian noise samples. Scale values were then estimated using the mlds function in the **MLDS** package [Knoblauch and Maloney, 2008] to fit the standard model. Thus, simulations generated under both the standard and diminishing-returns models were analyzed using the standard MLDS model to evaluate whether apparent diminishing returns could arise from estimation bias.

Each simulation condition was replicated 200 times. For each stimulus level, the mean estimated scale value across the 200 replications was used to characterize the average recovered scale. Unless otherwise indicated, intervals reported for parameter estimates are 95% Monte Carlo confidence intervals, defined by the 2.5th and 97.5th percentiles of the estimates obtained across repeated simulations. As only the shapes of the functions are of interest, each estimated scale was normalized by its scale value at the maximum stimulus level.

#### Sigmoid function

We also investigated a sigmoidal-shaped transducer function defined by the logistic function

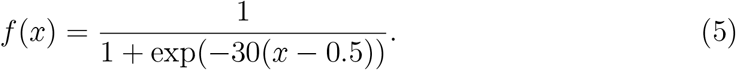

The scale was sampled at 10 stimulus levels on the interval (0, 1) either using a USS or URS sequence. The session length, thus, was reduced to 5 ∗ 120 = 600 triads. The number of repetitions was the same as that used when analyzing the power functions.

Simulations were performed for each of the four diminishing-returns exponents tested for power functions. All other procedures for fitting the standard MLDS model and estimating the average perceptual scale values and confidence intervals were unchanged.

The simulation framework was therefore designed to distinguish the effects of non-linear transduction of stimulus intensity from bias introduced by stimulus sampling, and to determine how these factors influence the recovery of perceptual geometry and its dependence on the diminishing-returns exponent.

### Assessment of parameter identifiability

To investigate the geometry of the likelihood surface, log likelihoods were calculated over a 200 × 200 grid of values of the transducer exponent, *p*, and diminishing-returns exponent, *α*. These likelihood-surface analyses were performed on individual simulated sessions rather than on averages across replications. Eight analyses were performed for the four combinations of *p* = (0.125, 8) and *α* = (0.5, 1), each evaluated with both USS and URS sequences. Contour plots were used to visualize the relative log-likelihood surfaces after subtracting the maximum log likelihood so that the peak value was zero (Supplmentary S5).

The local geometry of the likelihood surface around the grid location of the maximum likelihood was characterized by fitting the quadratic

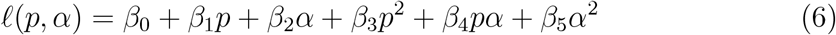

to a grid of 7 × 7 points centered at the maximum value of the surface. The local observed information matrix at the peak of the likelihood surface was approximated by

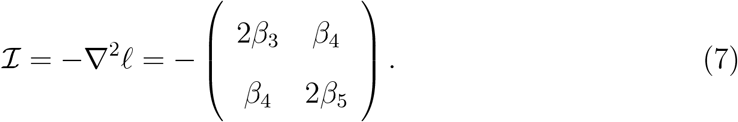

The eigenvalues of ℐ, (*λ*_1_, *λ*_2_), were used as measures of local curvature along the principal axes. The determinant of ℐ was calculated as the product of the eigenvalues to provide a measure of overall curvature. The condition number, *κ*, was calculated as the ratio of the largest to the smallest eigenvalue as an indication of anisotropy in the surface. The orientation of the principal axis (ridge direction), *θ*, was calculated as

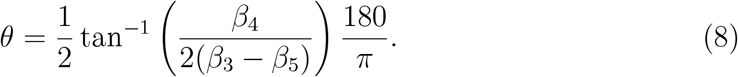

The robustness of these estimates was evaluated by repeating the analyses for windows of 11 × 11, 15 × 15, 21 × 21, 31 × 31 and 41 × 41 points around the maximum (Tables S1–S8). The estimates were highly stable across local window sizes, with little change in the determinant, condition number, or ridge angle over this range.

## Results

### Power transducer functions

#### Uniform Stimulus Sampling (USS)

The relative effects of *α* on the estimated scale shape with USS sequences are shown in Figure 2. The scales have been normalized by their maximum values in order to emphasize their differences in shape. Each panel shows data simulated with the transducer-function exponent indicated in the upper strip. Different point types show the mean estimated scale values and 95% confidence intervals for the four values of the diminishing-returns exponent, *α*, indicated in the legend. The solid curve in each panel shows the ground-truth power function used to simulate the data.

**Figure 2.**
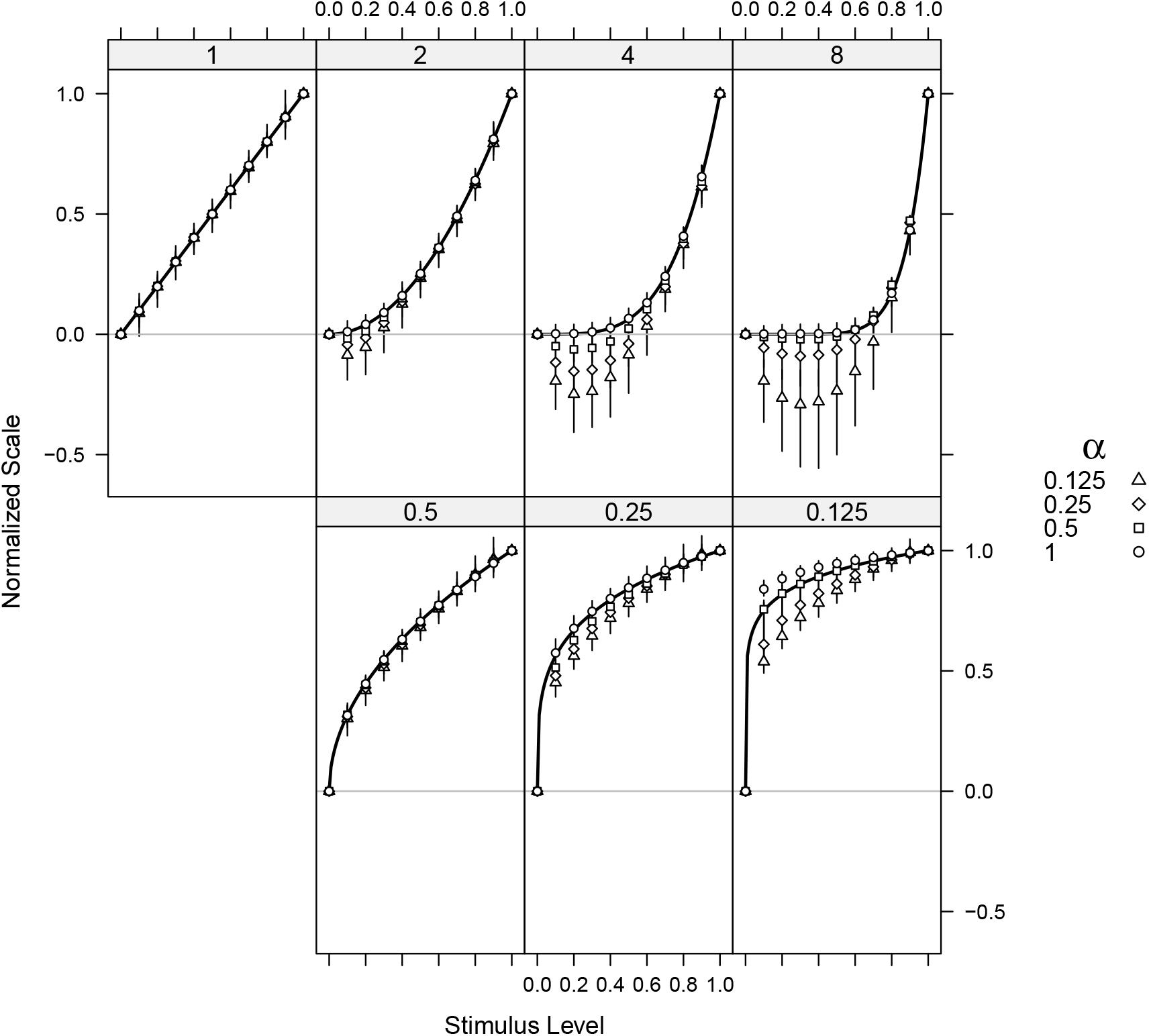
Normalized estimated perceptual scales for USS sequences. Each panel corresponds to data generated with a different transducer exponent, indicated in the upper strip. Four values of the diminishing-returns exponent using different point types are plotted in each panel. The solid curves are the ground-truth power functions for the data simulated in each panel. Error bars indicate 95% confidence intervals.

Remarkably, for the linear transducer (*p* = 1), the estimated scales fall almost exactly on the generating function, with negligible bias across values of the diminishing-returns exponent, *α*. For nonlinear transducers, however, the deviations depend on both the shape of the transducer and the value of *α*. For expansive transducers (*p >* 1), the estimates increasingly undershoot the generating function in its shallow regions. For strongly compressive transducers, as *p* decreases from 0.25 to 0.125, the deviations become more pronounced, with their magnitude and direction depending on *α* and the largest discrepancies occurring where the transducer changes most rapidly. Notably, for *p* = 0.125 and *α* = 1, the estimated scale slightly overestimates the underlying transducer. Together, these results show that the influence of the diminishing-returns exponent on the recovered scale depends on the degree and direction of transducer nonlinearity.

#### Uniform Response Sampling (URS)

The preceding results indicate that bias increases when stimulus density is mismatched to the rate of change of the transducer function. With expansive transducers (*p >* 1), the slope of the transducer is shallow at low stimulus intensities, and this region is over-sampled by USS, resulting in many samples in a region where the function changes little. Conversely, with compressive transducers (*p <* 1), the slope is steep at low stimulus intensities and this region is under-sampled by USS. The only condition showing unbiased results for all values of *α* is *p* = 1, for which uniform sampling in stimulus intensity is also uniform sampling in response. We therefore examined scale estimates when stimuli were sampled uniformly in response (URS; Figure 1b). If *S* is a USS stimulus sequence, then for power functions with exponent *p, S*^1*/p*^ generates a URS sequence.

Figure 3 shows the estimated normalized scale values when stimuli were selected according to a URS sequence and scale values were estimated using the standard decision rule. The solid curve shows the ground-truth transducer function used to generate the responses. In contrast to USS, URS samples the shallow regions of the transducer sparsely and the steep regions densely. For all values of *p*, the four estimated scales for different values of *α*, shown by the points, completely overlap. Thus, although the value of *α* used to generate the data differs by as much as a factor of four from the value used to estimate the scales, the estimated scales remain unchanged. The estimates are essentially unbiased, falling directly on the ground-truth transducer function (Figures S1 and S2). We suggest in the Discussion how such a URS sequence can be obtained.

**Figure 3.**
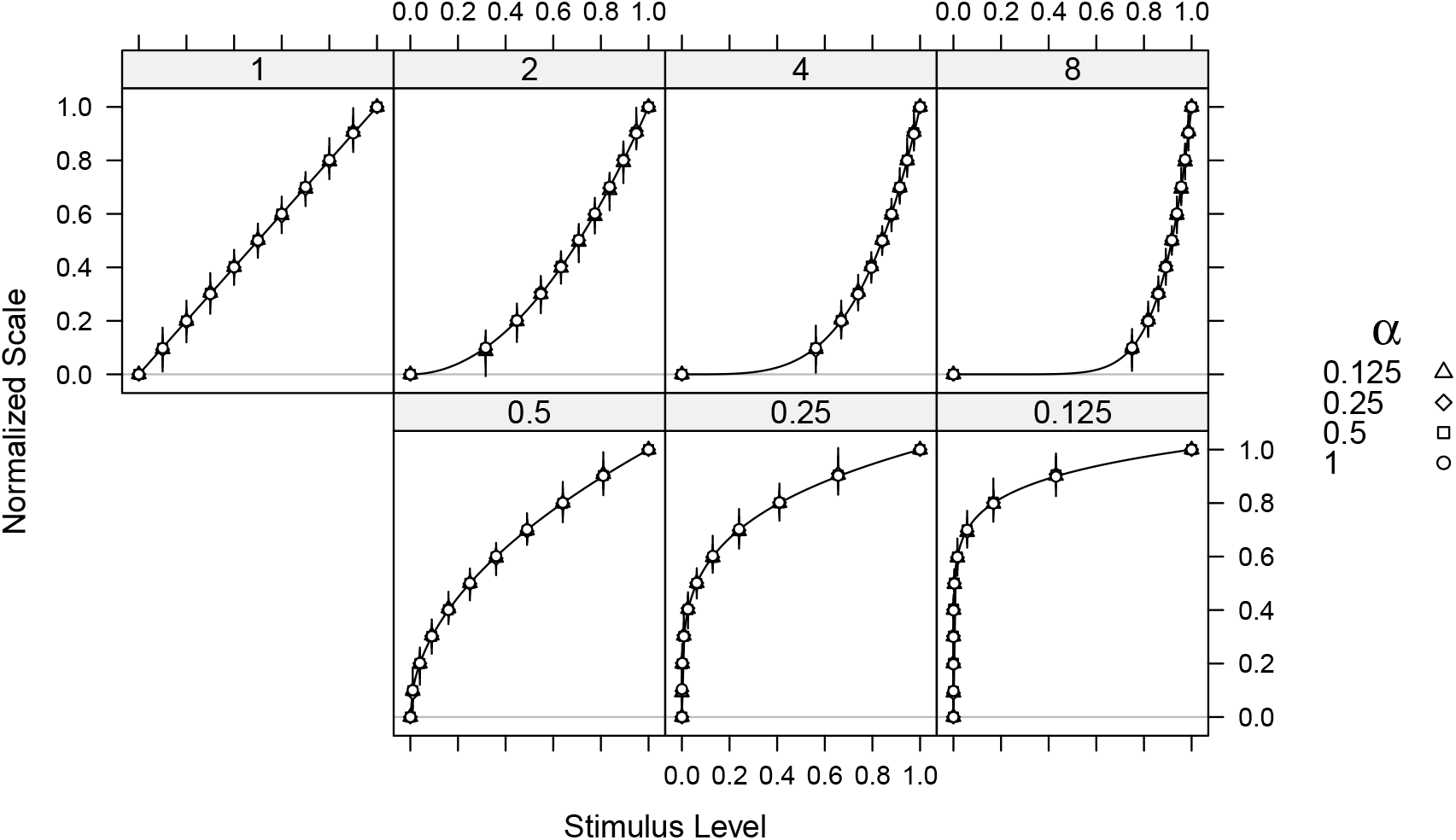
Normalized estimated perceptual scales for URS sequences. The format is the same as that used in Figure 2. Each panel corresponds to data generated with a different transducer function exponent, indicated in the upper strip. Four values of the diminishing-returns exponent, *α*, are plotted with different point types in each panel. The solid curves show the ground truth power functions for the data simulated in each panel. Error bars indicate 95% confidence intervals.

### Sigmoidal transducer function

Sigmoidal functions have regions of relatively flat and steep slope and, thus, if poorly sampled can show both over- and under-sampling effects. The left graph of Figure 4 demonstrates this for USS sampling using a logistic function (Equation 5). On the flat sections on the left and right of the scale where the function is over-sampled, there is increasing bias as the diminishing-returns exponent decreases. Notably, the two over-sampled tails exhibit biases of opposite sign. These results suggest that the form of the bias depends on the local structure of the transducer. On the rising portion of the curve where the function is under-sampled, the points are biased as well.

**Figure 4.**
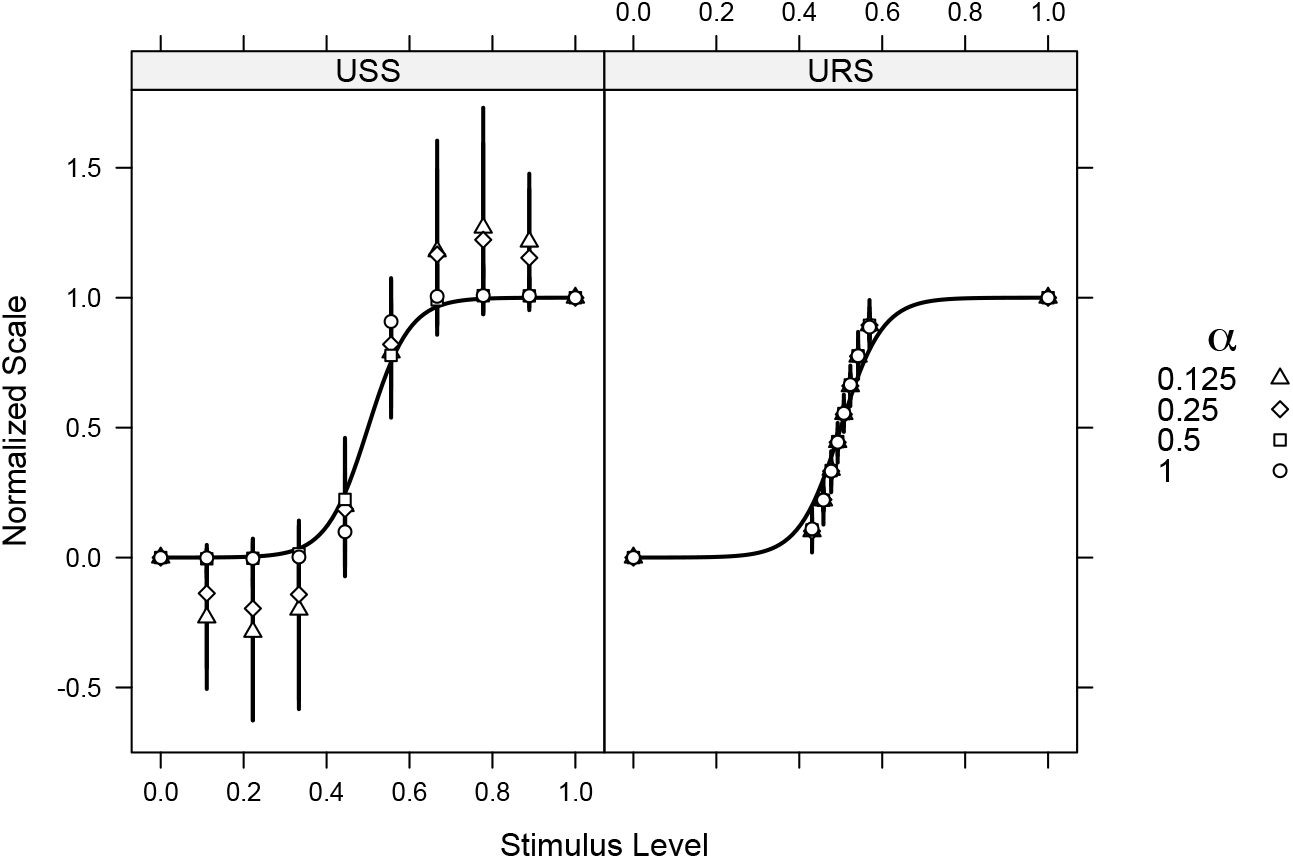
Normalized estimated perceptual scales for a sigmoidal transducer. The panels show data generated for each of the four values of diminishing-returns exponent, *α*, indicated by the point types in the legend to the right. The stimulus sequence, USS or URS, is indicated in the upper strip. The solid curves show the ground truth sigmoid function for the data simulated in each panel. Error bars indicate 95% confidence intervals.

In the right graph, the stimulus spacing has been adjusted to a URS sequence. Only one point is allocated to each of the flat regions and the rest are closely spaced on the rising portion. The results for all four values of *α* now overlap, as was obtained with power functions, indicating that the estimated scales are essentially independent of the diminishing-returns exponent. This stimulus strategy does not completely eliminate bias, however. As the points deviate from the central stimulus level, the averages are slightly above or below the ground-truth curve. These deviations are much smaller than those shown for the USS sequences.

The results of the above simulations raise two possibilities. The first is that *α* has little or no influence on the estimated scale and therefore need not be estimated. The second is that estimates of *α* are confounded with the slope of the transducer and sampling of the stimulus range, producing a ridge in the log-likelihood surface that is particularly pronounced when stimuli are poorly sampled. If the latter explanation is correct, optimally placed stimuli should reduce the correlation between the two parameters, even if *α* continues to have only a weak influence on the estimated scale values. We first consider why uniform response sampling produces this negligible influence of *α* on the estimated scale, and then examine the local geometry of the log-likelihood surface for power functions to determine whether *α* remains confounded with the transducer slope.

### Why uniform response sampling removes the effect of diminishing returns

The invariance to *α* under URS can be understood directly from the structure of the decision rule. We develop the argument explicitly for the power-law transducer considered here; the same reasoning applies more generally to any monotonic transducer sampled uniformly in response space. Consider first the special case of a linear transducer (*p* = 1). Ignoring the noise term, the generalized decision variable for triads is

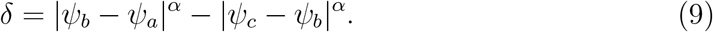

Since *ψ*(*x*) = *x*, the stimulus set is fixed and equally spaced under both USS and URS sequences. Let the stimulus levels be *x*_*i*_ = *i*Δ*x*. Every interval difference is then

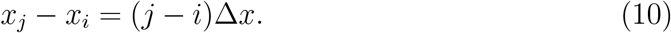

and for arbitrary stimulus indices *i, j, k*, the generalized difference can be written

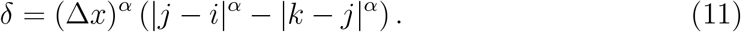

The factor (Δ*x*)^*α*^ is a common multiplicative constant and can therefore be absorbed into the noise scale, while the remaining *d*^*α*^ transformation is not a common scaling factor across trials. The ordinal information in the comparisons consequently depends only on the integer interval lengths raised to the power *α*. Because the MLDS scale values at the *q* stimulus levels are free parameters, the model seeks a set of scale values, 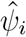, that reproduces the ordering of the observed interval comparisons.

For uniformly spaced levels, the transformation *d* → *d*^*α*^ preserves the ordering of interval sizes for every *α >* 0.

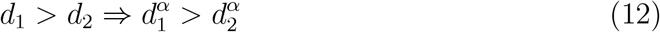

for *α >* 0 and *d* = |*ψ*(*x*_*j*_) − *ψ*(*x*_*i*_)|. Thus, changing *α* changes the magnitude of the decision variable but not the rank ordering of the interval comparisons. The estimated MLDS scale therefore has the same normalized shape, apart from an arbitrary scale factor. The situation changes when

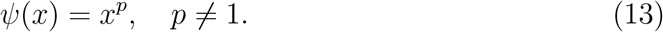

The intervals in perceptual response space are then

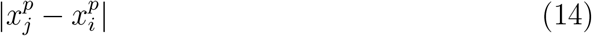

so equal stimulus steps no longer correspond to equal response-space intervals. Changing *p* therefore changes the relative sizes of the intervals, and applying the exponent *α* can alter the relative differences between intervals.. The MLDS scale can consequently depend on *α*, producing the bias observed with USS.

URS, however, restores equal spacing in response space, the condition underlying the *p* = 1 result. Rather than spacing stimuli uniformly in the physical stimulus domain, the stimuli are chosen so that their transducer responses are equally spaced.

That is, the stimulus levels are selected to satisfy

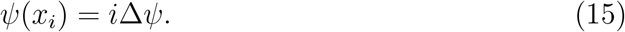

This condition is sufficient for the argument below and does not depend on the particular form of the transducer. For the power-law transducer used in the simulations, it gives

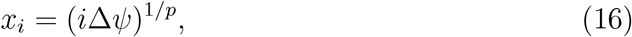

and hence

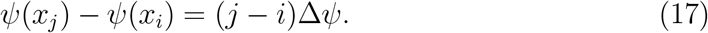

Thus, regardless of the value of *p*, the sampled points are equally spaced in response space. The interval structure is therefore identical to that of the linear case, and the argument above applies: raising the interval lengths to the power *α* changes their magnitudes but preserves their ordering. Under URS, *α* does not change the ordering of noiseless interval comparisons. The same argument applies to quads.

### Log Likelihood Surfaces

We next examined the local geometry of the log-likelihood surface for power functions to determine whether uniform response sampling also reduces the confounding between *α* and the transducer slope. Likelihoods were calculated over a grid of values of *p* and *α* for data sets generated with fixed values of these parameters. Contours were plotted at specified decreases in log likelihood from the maximum. Data sets were generated for the four combinations of *p* = 0.125 and 8 and *α* = 0.5 and 1, using both USS and URS stimulus sequences. These extreme parameter combinations were chosen to emphasize the effect of stimulus sampling on the geometry of the likelihood surface under strongly nonlinear transduction and substantial differences in the scaling of perceptual intervals. For ease of comparison, the coordinate axes are matched across each pair of graphs generated with the same (*p, α*) combination. This likelihood analysis is not intended as an alternative estimation procedure for MLDS. Rather, it serves as a diagnostic tool for understanding the geometry of the likelihood under a two-parameter transducer model. By parameterizing the model in terms of two interpretable parameters—the transducer exponent and the diminishing-returns exponent—we can visualize parameter identifiability and the extent to which the sampling design induces confounding between parameters *p* and *α*.

#### Compressive Transducer Functions

Figure 5a and b show contours of equal log likelihood for data generated with *p* = 0.125 and *α* = 1 using USS and URS sequences, respectively. The color gradients in each figure reflect the change in likelihood across the parameter space varying from white to a deep orange as the likelihood decreases from the maximum. The likelihood at the true parameter values is indicated by the black circle in both panels (and in subsequent plots). The maximum does not occur exactly at the true parameter values, as expected for a single simulated session, but lies reasonably close to them. These surfaces illustrate the local geometry for individual sessions rather than providing estimates of sampling bias.

**Figure 5.**
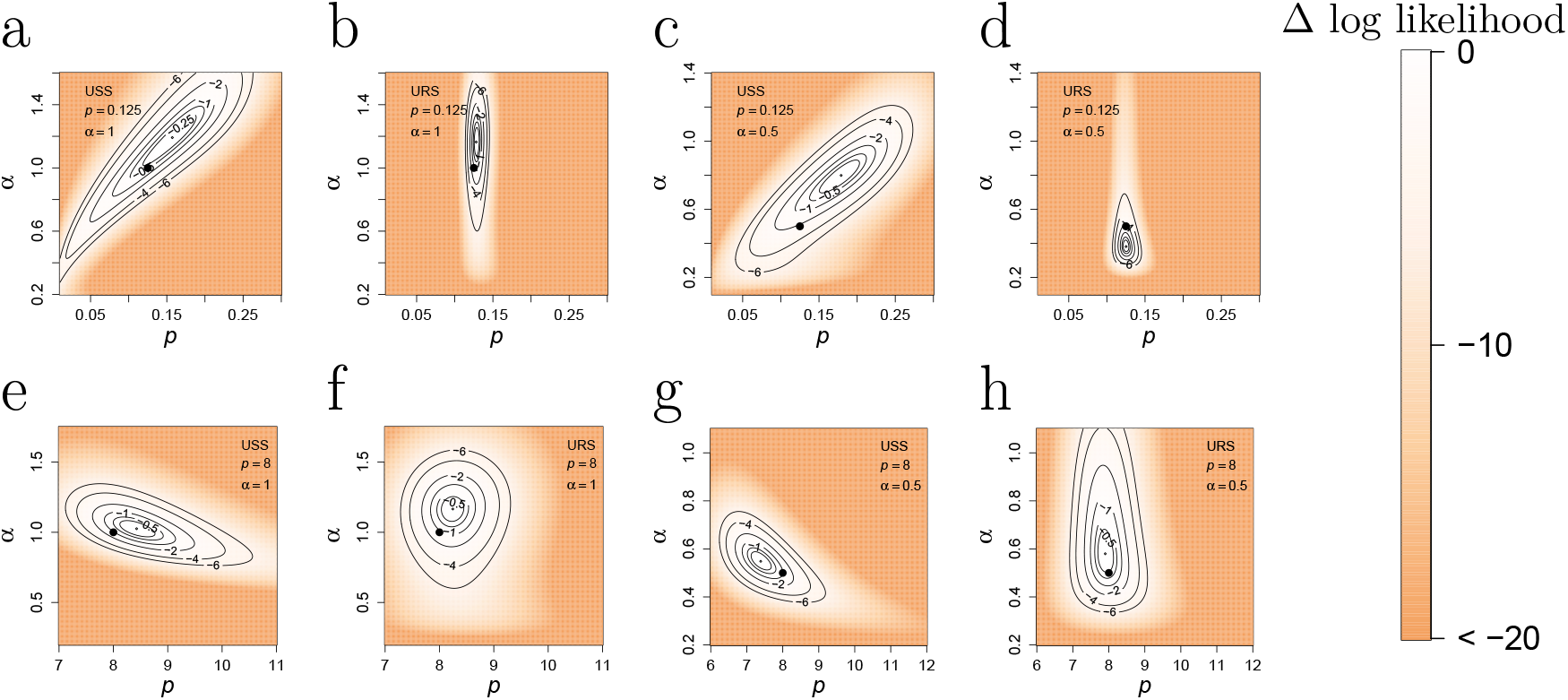
Relative log-likelihood surfaces for several (*p, α*) combinations. The parameter values used to generate each surface are indicated by the black circle. Panels a, c, e, and g show surfaces generated using USS sequences; panels b, d, f, and h show the corresponding surfaces generated using URS sequences. The (*p, α*) pair and the stimulus sequence for the surface are indicated within each panel. Contours correspond to decreases in log likelihood from the maximum of -0.5, -1, -2, -4, and -6. The orange color gradient in each image shows the smooth change in log likelihood as indicated by the color bar on the right.

The large condition numbers, *κ*, (Table 1) indicate that the local likelihood surfaces are strongly anisotropic, with one principal direction substantially less identifiable than the other. Notably, for USS (Figure 5a), the elongated contours are oriented approximately along the *p* = *α* direction, indicating a strong trade-off between the two parameters. This is consistent with the ridge angle, *θ*, which is close to 45 deg. The approximately 50-fold larger determinant for URS (Figure 5b) indicates substantially greater overall local curvature, with the contours correspondingly more localized. Their elongation is now approximately parallel to the *α* axis (*θ* close to 90 deg), and the small eigenvalue in this direction is only about 3% of the larger one, indicating that *α* remains poorly identifiable even with URS sampling. These qualitative patterns are preserved when the likelihood surface is examined in (log *p*, log *α*) coordinates. A similar pattern emerges in Figures 5c and d for the parameter combination (*p, α*) = (0.125, 0.5), although *κ* decreases by only a factor of 3.5 for the URS sequences.

**Table 1:** Parameter estimates to characterize local geometry of the log likelihood surfaces. Columns 1–2, values of *p* and *α* to generate the data set; Column 3, stimulus-spacing regime (USS or URS), Column 4–5, maximum-likelihood estimates of the transducer and diminishing-returns exponents, respectively; Columns 6–7, eigenvalues of the observed information matrix *I*, representing the local curvatures of the log-likelihood surface; Column 8, determinant of *I*; Column 9, condition number, calculated as the ratio of the larger to smaller eigenvalue of *I*; Column 10, ridge angle, defined as the orientation of the shallow direction of the likelihood surface around its maximum.

| p | α | Sampling | p̂ | α̂ | λ<sub>1</sub> | λ<sub>2</sub> | det(ℐ) | κ | θ (deg) |
|---|---|---|---|---|---|---|---|---|---|
| 0.125 | 1.0 | USS | 0.157 | 1.192 | 0.013 | 4.62e-04 | 6.08e-06 | 28.500 | 43.3 |
| 0.125 | 0.5 | USS | 0.179 | 0.799 | 0.012 | 0.001 | 1.23e-05 | 12.300 | 38.2 |
| 8.000 | 1.0 | USS | 8.427 | 1.026 | 0.015 | 0.002 | 2.67e-05 | 8.400 | 162.6 |
| 8.000 | 0.5 | USS | 7.387 | 0.548 | 0.016 | 0.004 | 6.45e-05 | 4.000 | 139.9 |
| 0.125 | 1.0 | URS | 0.128 | 1.164 | 0.104 | 0.003 | 3.3e-04 | 32.900 | 89.4 |
| 0.125 | 0.5 | URS | 0.125 | 0.381 | 0.073 | 0.019 | 0.001 | 3.800 | 94.5 |
| 8.000 | 1.0 | URS | 8.246 | 1.166 | 0.005 | 0.004 | 1.85e-05 | 1.300 | 74.0 |
| 8.000 | 0.5 | URS | 7.899 | 0.580 | 0.010 | 0.001 | 1.29e-05 | 7.600 | 93.5 |

#### Expansive Transducer Functions

When the transducer function is expansive, over-sampling occurs over a range where the function changes gradually rather than under-sampling the region where it changes rapidly. Log-likelihood surfaces for this case are shown in Figures 5e–h for both USS and URS stimulus sequences. As in the compressive condition, the USS sequences (Figures 5e and g) generate a ridge in the likelihood surface, as indicated by the large values of *κ*, demonstrating a *p*–*α* trade-off in estimation. The orientation is reversed, however, with *θ* near 150 deg, indicating a negative association between the two parameters. With URS sampling (Figures 5f and h), this trade-off is sub-stantially reduced with *θ* near 90 deg. The condition number, *κ*, is reduced to nearly unity for the (8, 1) condition, indicating that the local curvature is approximately isotropic. The eigenvalues are also more similar in magnitude, indicating that neither parameter is substantially less identifiable than the other.

Consistent with the scale estimates, the maximum-likelihood estimates of *p* and *α* show more bias under USS sequences than for URS. Across the four parameter combinations examined, URS generally rotated the likelihood ridge away from a *p* − *α* trade-off and toward the *α* direction. The condition number was reduced for two of the four combinations, but increased for the other two, indicating that URS does not uniformly improve overall isotropy. The particularly striking case is (8, 1) where *κ* decreases from 8.5 to 1.3. URS consistently reduced the *p* − *α* confounding, as evidenced by rotation of the likelihood ridge toward the *α* axis. This reduction in confounding did not always coincide with improved local isotropy of the likelihood surface though, indicating that parameter identifiability also depends on the intrinsic geometry of the model. Although the surfaces shown in Figure 5 are derived from individual simulated sessions and therefore vary slightly across repetitions, the main qualitative features are highly reproducible: USS consistently produces likelihood ridges aligned with a *p* − *α* tradeoff, whereas URS rotates the ridge toward the *α* axis and typically increases local curvature around the optimum. Thus, the precise contours vary between repetitions, but the geometric signatures associated with stimulus sampling remain stable

## Discussion

### Sampling and the geometry of finite intervals

The dependence of psychophysical parameter estimation on stimulus sampling has been recognized previously. Wichmann and Hill [2001], for example, demonstrated that the placement of stimulus samples can substantially affect the bias and precision of estimates of psychometric-function parameters. Our results reveal a related but distinct consequence of stimulus sampling for MLDS: sampling the physical stimulus domain uniformly can bias the recovered perceptual scale when the physical-to-perceptual transducer is nonlinear. MLDS does not probe infinitesimal geometry directly; rather, it infers geometry from finite intervals through ordinal judgments. As a result, the recovered geometry is potentially dependent on how the stimulus continuum is sampled. With a nonlinear transducer, uniform sampling in the physical stimulus domain produces nonuniform sampling in perceptual space, leading to local under-sampling or over-sampling. These sampling effects can systematically interact with the rules for composing finite intervals in the representation being compared, introducing bias into the estimated geometry.

The usual requirement that the stimulus set satisfy an ordering property provides some protection against one aspect of this sampling problem. In MLDS, stimulus levels should be sufficiently well separated for observers to order them reliably along the stimulus continuum [Maloney and Yang, 2003, Knoblauch and Maloney, 2012, Maloney and Knoblauch, 2020, Teti et al., 2022]. This requirement should discourage over-sampling of regions in which the transducer changes slowly, because stimuli that are too closely spaced may become difficult to order reliably. It does not, however, protect against under-sampling regions in which the transducer changes rapidly: stimuli separated by large perceptual differences can remain readily orderable even when too few levels are available to characterize the local geometry. A warning about GLM fitted probabilities numerically equal to 0 or 1, such as that issued by the R GLM implementation, may provide an additional indication that some intervals are very large relative to the noise in perceptual space. Such warnings are not, however, specific to under-sampling: we observed them systematically for particular combinations of *α* and *p*, including *p >* 1 (Figure S3). Their occurrence should therefore be treated as a diagnostic cue rather than as evidence of under-sampling, prompting examination of the spacing of the stimulus levels in relation to the estimated scale [Knoblauch and Maloney, 2012].

The ordering principle also places a qualification on the construction of a URS sequence. For a monotonic transducer, ordering the stimuli by their physical level also orders them along the perceptual scale, and therefore, uniformly spaced perceptual levels can be mapped back to stimulus levels using the inverse transducer. For a nonmonotonic transducer, however, physical stimulus order need not correspond to perceptual order: increasing the stimulus level may initially increase the perceptual response and then decrease it, such that a higher stimulus level can have a perceptual value intermediate between those of lower stimulus levels. In this case, no single global inverse transducer yields a physically ordered URS sequence that satisfies the ordering prerequisite. URS sequences could instead be constructed separately over monotonic portions of the transducer. Importantly, this limitation concerns the construction of the stimulus sequence, not MLDS estimation itself, which does not require the transducer to have a specified parametric form.

We examined bias in MLDS scale estimation across a range of transducer non-linearities, perceptual-space geometries, and stimulus-sampling schemes. With uniform sampling in the stimulus domain and no diminishing-returns transformation, MLDS was robust except under the most extreme compressive nonlinearities. When diminishing-returns behavior was incorporated into the model but not accounted for in estimation, however, systematic bias emerged for nonlinear transducers. The bias arose from two complementary sampling effects: under-sampling where the transducer changed rapidly and over-sampling where it changed gradually. Both effects were substantially reduced when stimuli were sampled uniformly in response space. Importantly, under uniform response sampling, scale estimates were essentially invariant to the diminishing-returns transformation.

Although the principal simulations used power functions, the mechanism identified here is not specific to that family. Power functions provide a convenient family of transducers spanning a wide range of compressive and expansive nonlinearities and, importantly, a range of local slopes. In addition, we demonstrated under- and over-sampling within a single function using a sigmoid-shaped transducer that displays both relatively flat and steep regions. The sampling effects identified here arise from the nonuniform representation of perceptual space produced by a nonlinear trans-ducer, rather than from any specific property of the power function. More generally, the relevant quantity is not the functional form itself but the local rate at which the transducer maps physical stimulus intervals into perceptual space. Any smooth nonlinear transducer with regions of relatively rapid and gradual change, such as the logistic function used here, will therefore produce analogous under-sampling and over-sampling effects under uniform physical sampling.

Importantly, under uniform response sampling, scale estimates were essentially unchanged across the diminishing-returns transformations. The mathematical result clarifies why the recovered MLDS scale is invariant to *α* under URS. By sampling uniformly in response space, URS makes the interval comparisons depend only on the ordinal positions of the sampled stimuli. The *α* transformation therefore changes the magnitude of the decision variable without changing the ordering of noiseless comparisons. The simulations show that, at the noise level considered here, this invariance translates into essentially identical recovered scales across values of *α*.

The likelihood surfaces provide a complementary view of these results. The geometry of the likelihood depended strongly on stimulus spacing, with uniform response sampling generally reducing confounding between the transducer and diminishing-returns exponents. Under USS, the shallow direction of the likelihood surface was oriented along a *p* − *α* trade-off, whereas under URS it was generally oriented more nearly along the *α* axis. Thus, URS reduced the extent to which transducer nonlin-earity and diminishing returns could substitute for one another in explaining the data. It did not, however, uniformly improve parameter identifiability: *α* remained weakly identified in some conditions, whereas the likelihood surface was nearly isotropic for *p* = 8, *α* = 1.

Although the estimated perceptual scales were largely insensitive to *α* under URS, the likelihood surfaces obtained under the restricted power-function parameterization retained some curvature in *α*. These results are not contradictory. The semi-parametric MLDS fit estimates the scale at each stimulus level without imposing a functional form, whereas the likelihood surfaces restrict the scale to the two-parameter family *ψ*(*x*) = *x*^*p*^. Thus, the curvature in *α* reflects the distinguishability of *p* and *α* within this restricted parametric model and does not imply that *α* materially changes the recovered perceptual scale. In this sense, the semi-parametric fits provide the more direct evidence that the estimated scale is insensitive to *α* under URS, whereas the likelihood surfaces reveal residual parameter confounding within the parametric model.

### Implications for previous and future MLDS studies

Our findings relate to recent work examining sampling in MLDS. Aguilar et al. [2023] investigated whether subsets of the full set of possible triads could be used to reduce the number of trials required for scale estimation. Their simulations showed that the accuracy of the estimated scale depended on the sampling scheme and, in particular, that subsampling could introduce bias when the underlying perceptual scale was nonlinear. They proposed selecting triads according to the difference between the stimulus indices, thereby preferentially sampling comparisons expected to provide useful information about the scale. Our results identify a complementary source of sampling bias: even when all triads are retained, uniformly spacing stimuli in the physical stimulus domain can systematically bias estimation when the transducer is nonlinear. Thus, both the selection of triads and the spacing of stimulus levels can affect the accuracy of MLDS estimates, although through different mechanisms.

These two sampling issues represent distinct aspects of experimental design: which stimulus levels are presented and which comparisons among those levels are made. The present simulations isolate the former by including all possible triads, showing that bias can arise from stimulus spacing alone.

The implications of these sampling effects extend to previous applications of MLDS. Some published studies have used sampling schemes that could be susceptible to the biases identified here. Potential over-sampling of relatively flat regions of the transducer function occurs in studies using MLDS to investigate the levels of compression that produce perceptual distortion [Charrier et al., 2007], the perception of correlation in scatterplots [Knoblauch and Maloney, 2012] and slant perception [Aguilar et al., 2017]. The latter study shows a pattern that resembles the characteristic undershoot of the transducer function illustrated above. Conversely, the pilot MLDS estimates for the “red chroma” condition in Rogers et al. [2016] can be interpreted in light of potential under-sampling: the stimuli were equally spaced on the chroma axis, but the estimated scale shows a steep increase between the first two stimulus levels.

A separate issue concerns nonadditivity in the contribution of perceived intervals to the judgment, which can produce diminishing returns. Teti et al. [2022] provide an important example: the relationship between the magnitude of a perceptual difference and its contribution to the judgment may itself be nonlinear. These are conceptually distinct sources of nonlinearity.

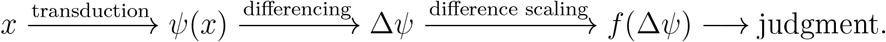

Nonlinearity can occur both in the transduction from stimulus to perceptual magnitude, indicated by the first arrow above, and in the transformation of perceptual differences into decision evidence, indicated by the third arrow. However, these two sources of nonlinearity need not have equivalent consequences for MLDS. Our simulations show that even when *α* ≠ 1, appropriate sampling in perceptual-space can permit recovery of the underlying perceptual scale with little bias. Therefore, future MLDS experiments should distinguish two questions: What is the perceptual geometry, and how are perceptual differences transformed into decision evidence? Importantly, sampling should be designed with the first question in mind, rather than assuming that the physical stimulus domain is neutral.

If the primary goal of MLDS is to estimate the transducer function, this raises a practical question: uniform response sampling requires some prior knowledge of the approximate shape of that function. In practice, this information could be obtained from pilot experiments used to estimate the perceptual scale. The objective is to select stimulus levels that yield approximately equal increments in the estimated transducer output. Equivalently, the transducer function should be approximately linear when expressed as a function of the ordinal ranks of the selected stimulus levels. This formulation also provides a convenient basis for estimating transducer functions with generalized linear mixed models (GLMMs) when data from multiple observers are analyzed jointly. Thus, response-space sampling should be viewed as an iterative design strategy rather than as a requirement for prior knowledge of the exact transducer. Perfectly uniform response sampling may not be necessary; rather, the goal is to allocate more stimuli to regions where the transducer changes rapidly and fewer to regions where it changes gradually. In some cases, a simple nonlinear sampling scheme, such as logarithmic spacing, may provide an adequate approximation to uniform response sampling [Bellot et al., 2016, Knoblauch et al., 2020, Shooner and Mullen, 2022].

These results suggest two related uses of response-space sampling. When the primary goal is to estimate the transducer function, approximately uniform sampling in perceptual space can reduce the influence of the decision rule on the recovered scale. When the goal is to characterize perceptual geometry, comparisons between scales estimated under USS and URS may help reveal the influence of the decision rule on the estimated scale. The extent to which such comparisons can reliably identify departures from the assumption that *α* = 1 remains to be investigated.

Because the decision models are closely related, we expect these results to be relevant to Maximum Likelihood Conjoint Measurement (MLCM) studies as well. URS-like stimulus sequences have been suggested for MLDS [Knoblauch and Maloney, 2012] and used in several MLCM studies [Lisi and Gorea, 2016, Rogers et al., 2016, 2018, Abbatecola et al., 2021, 2025], where linear or quadratic parametric representations of the perceptual scales reduced the number of free parameters and facilitated the use of generalized linear mixed-effects models.

### Implications for perceptual geometry

The results demonstrate that the observability of departures from a proposed perceptual geometry can depend critically on stimulus sampling. An observer with an additive decision rule can exhibit apparent non-Riemannian behavior when stimulus sampling inadequately represents the perceptual scale. Both under-sampling regions where the transducer changes rapidly and over-sampling regions where it changes gradually can distort the estimated scale. When stimuli are sampled approximately uniformly in perceptual space, these apparent deviations are substantially reduced. More generally, apparent departures from a proposed perceptual geometry should be interpreted with caution unless the experimental design adequately represents the space in which perceptual differences are evaluated.

This principle extends beyond MLDS: whenever a nonlinear transformation intervenes between the physical stimulus and the representation being modeled, nonuniform sampling can distort the estimated representation in ways that may be mistaken for departures from the assumed geometry. This concern is relevant to theoretical approaches that relate measured perceptual scales to perceptual geometry, including those based on Fisher information [Vacher and Mamassian, 2024]. Interpreting estimated perceptual geometry therefore requires distinguishing properties of the un-derlying representation from effects of stimulus sampling and the decision rule used to evaluate perceptual differences.

## Data availability

The code used to generate the simulations is provided in an R Markdown file. The simulations use a random seed that changes between runs, so the numerical results will vary slightly across runs, while the qualitative results remain unchanged. The R Markdown file will be made available in a public repository upon acceptance.

## Author Contributions

Conceptualizetion: K.K. ; Writing – original draft: K.K.; Writing – review and editing: C.A., K.B., K.K.; Data analysis: C.A., K.B., K.K. K.K. is the corresponding author and acts as the guarantor of the work, taking responsibility for the integrity of the paper as a whole. (ORCID iD: 0000-0002-4681-4638).

## Acknowledgments

We thank David Brainard and Pascal Mamassian for helpful discussion and suggestions about this work.

