## Supplemental Figures S1-S3 and Tables S1-S8 for "Stimulus Sampling Shapes Estimated Perceptual Geometry in Psychophysical Scaling"

#### This PDF file includes:

Figs. S1 to S3
Tables S1 to S8

### Evaluation of bias

The bias between the average simulated results and ground truth functions is plotted as a function of stimulus level for each $(p, \alpha)$ pair.

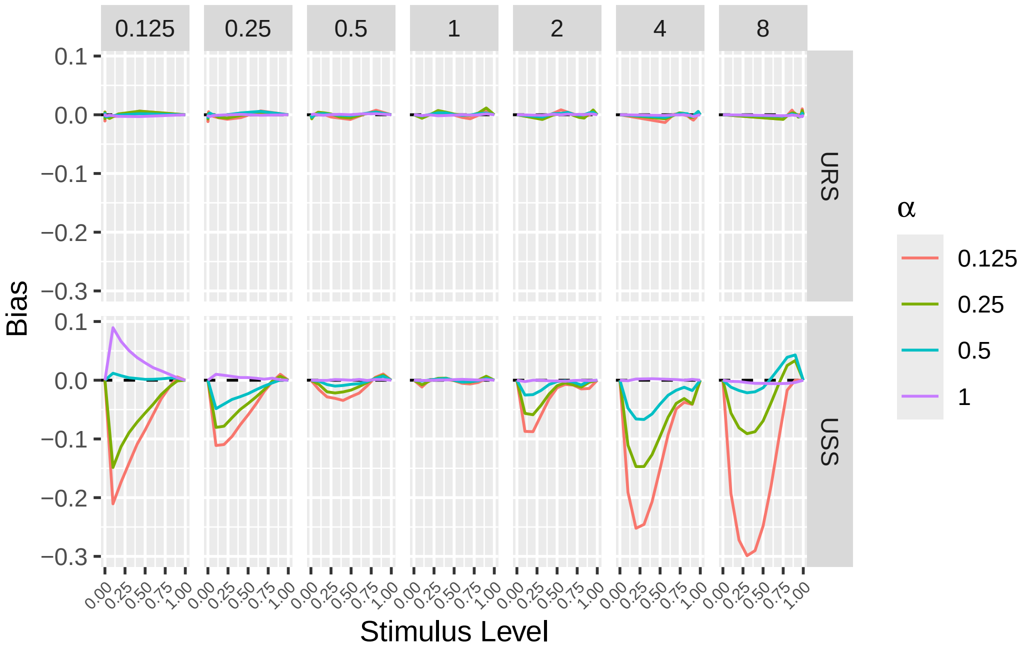


**Figure S1.** Bias, defined as the difference between the simulated function and the ground truth used as a generator in the simulations as a function of $\alpha$ with each panel corresponding to a different power exponent $p$. The top row is for 'URS' sequences and the bottom for 'USS'.

For each power, $p$, the root mean square error between the average simulated results and the ground truth function is plotted as a function of the diminishing returns exponent, $\alpha$.

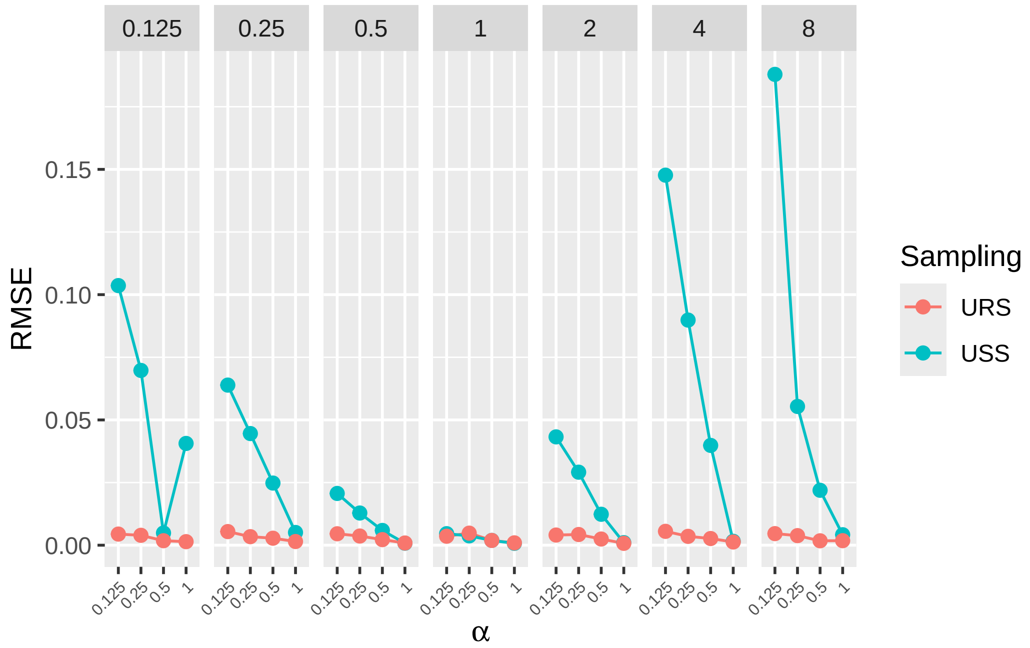


**Figure S2.** Root Mean Square Error (RMSE) plotted as a function of $\alpha$ for the two sampling sequences, as indicated in the legend. Each panel corresponds to a different power law exponent, $p$.

#### Distribution of warnings

The distributions of warnings are plotted for all of the $(p, \alpha)$ combinations.

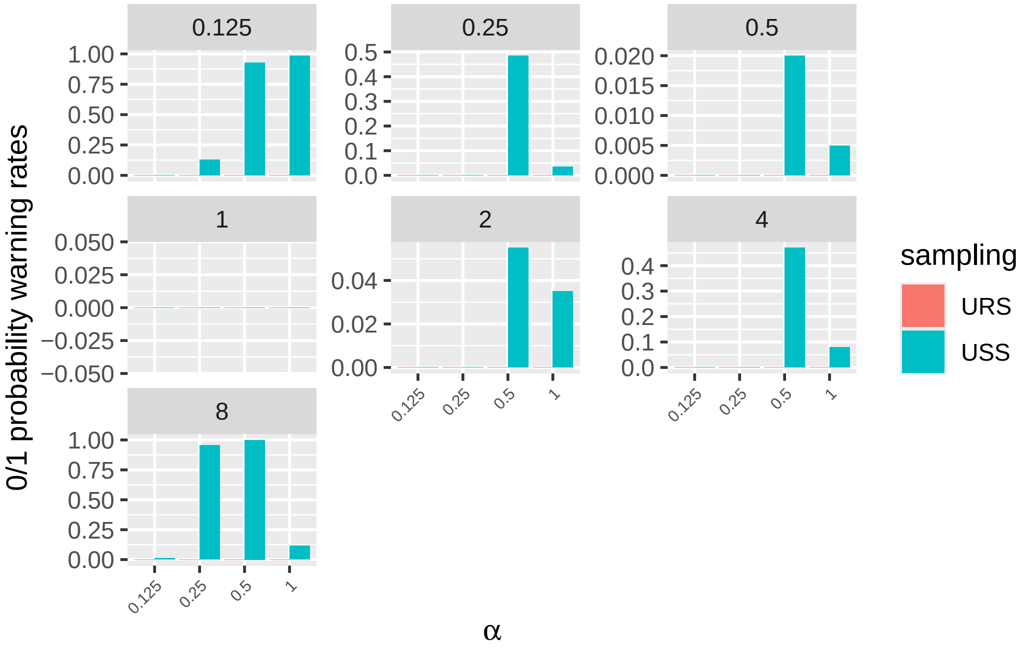


**Figure S3.** The proportion of warnings generated during the simulation for glm calls, indicating 0/1 probabilities estimated in the data. Each panel corresponds to a different power law exponent, $p$. The bars indicate the relative frequency of warnings as a function of $\alpha$. The bar colors indicate 'USS' or 'URS' sequences, although 'URS' sequences generated no warnings.

### Stability of local geometry around likelihood surface maximum

The stability of the estimations of the characteristics of the log likelihood surface around its peak is evaluated for each of the $p - \alpha$ pairs and sampling sequences displayed in Figure 5 of the article. Here, the determinant of the information matrix $\det \mathcal{I}$, the condition number $\kappa$ and the angle of the ridge $\theta$ are recalculated for grids around the maximum of $7 \times 7$, $11 \times 11$, $15 \times 15$, $21 \times 21$ and $41 \times 41$. For each combination of $p - \alpha$ and sampling sequence, the results are indicated in a table below, indicating how little the parameter estimates change with the size of the grid. The left column displays results for USS sequences and the table for the matching URS condition is displayed in the right column on the same row.

**Table S1**

| p | $\alpha$ | Sampling | resolution | $\det(\mathcal{I})$ | $\kappa$ | $\theta$ |
|---|---|---|---|---|---|---|
| 0.125 | 1.0 | USS | 7 x 7 | 1.887E-05 | 8.031 | 41.4 |
| 0.125 | 1.0 | USS | 11 x 11 | 1.888E-05 | 8.023 | 41.4 |
| 0.125 | 1.0 | USS | 15 x 15 | 1.889E-05 | 8.012 | 41.4 |
| 0.125 | 1.0 | USS | 21 x 21 | 1.890E-05 | 7.988 | 41.4 |
| 0.125 | 1.0 | USS | 41 x 41 | 1.898E-05 | 7.855 | 41.5 |

**Table S2**

| p | $\alpha$ | Sampling | resolution | $\det(\mathcal{I})$ | $\kappa$ | $\theta$ |
|---|---|---|---|---|---|---|
| 0.125 | 1.0 | URS | 7 x 7 | 2.584E-04 | 24.563 | 91.5 |
| 0.125 | 1.0 | URS | 11 x 11 | 2.599E-04 | 24.468 | 91.4 |
| 0.125 | 1.0 | URS | 15 x 15 | 2.621E-04 | 24.331 | 91.4 |
| 0.125 | 1.0 | URS | 21 x 21 | 2.667E-04 | 24.047 | 91.3 |
| 0.125 | 1.0 | URS | 41 x 41 | 2.957E-04 | 22.444 | 90.9 |

**Table S3**

| p | $\alpha$ | Sampling | resolution | $\det(\mathcal{I})$ | $\kappa$ | $\theta$ |
|---|---|---|---|---|---|---|
| 0.125 | 0.5 | USS | 7 x 7 | 6.321E-06 | 10.185 | 43.8 |
| 0.125 | 0.5 | USS | 11 x 11 | 6.376E-06 | 10.133 | 43.9 |
| 0.125 | 0.5 | USS | 15 x 15 | 6.456E-06 | 10.058 | 43.9 |
| 0.125 | 0.5 | USS | 21 x 21 | 6.629E-06 | 9.902 | 43.9 |
| 0.125 | 0.5 | USS | 41 x 41 | 7.789E-06 | 9.007 | 43.9 |

**Table S4**

| p | $\alpha$ | Sampling | resolution | $\det(\mathcal{I})$ | $\kappa$ | $\theta$ |
|---|---|---|---|---|---|---|
| 0.125 | 0.5 | URS | 7 x 7 | 5.181E-04 | 20.660 | 92.0 |
| 0.125 | 0.5 | URS | 11 x 11 | 5.193E-04 | 20.663 | 91.9 |
| 0.125 | 0.5 | URS | 15 x 15 | 5.210E-04 | 20.667 | 91.9 |
| 0.125 | 0.5 | URS | 21 x 21 | 5.247E-04 | 20.674 | 91.8 |
| 0.125 | 0.5 | URS | 41 x 41 | 5.473E-04 | 20.711 | 91.2 |

**Table S5**

| p | $\alpha$ | Sampling | resolution | $\det(\mathcal{I})$ | $\kappa$ | $\theta$ |
|---|---|---|---|---|---|---|
| 8.000 | 1.0 | USS | 7 x 7 | 4.990E-05 | 5.641 | 159.6 |
| 8.000 | 1.0 | USS | 11 x 11 | 4.997E-05 | 5.644 | 159.6 |
| 8.000 | 1.0 | USS | 15 x 15 | 5.007E-05 | 5.648 | 159.6 |
| 8.000 | 1.0 | USS | 21 x 21 | 5.026E-05 | 5.657 | 159.7 |
| 8.000 | 1.0 | USS | 41 x 41 | 5.141E-05 | 5.709 | 160.0 |

**Table S6**

| p | $\alpha$ | Sampling | resolution | $\det(\mathcal{I})$ | $\kappa$ | $\theta$ |
|---|---|---|---|---|---|---|
| 8.000 | 1.0 | URS | 7 x 7 | 1.685E-05 | 1.247 | 77.1 |
| 8.000 | 1.0 | URS | 11 x 11 | 1.683E-05 | 1.248 | 77.2 |
| 8.000 | 1.0 | URS | 15 x 15 | 1.681E-05 | 1.249 | 77.2 |
| 8.000 | 1.0 | URS | 21 x 21 | 1.677E-05 | 1.252 | 77.4 |
| 8.000 | 1.0 | URS | 41 x 41 | 1.653E-05 | 1.267 | 78.0 |

**Table S7**

| p | $\alpha$ | Sampling | resolution | $\det(\mathcal{I})$ | $\kappa$ | $\theta$ |
|---|---|---|---|---|---|---|
| 8.000 | 0.5 | USS | 7 x 7 | 5.537E-05 | 6.080 | 148.9 |
| 8.000 | 0.5 | USS | 11 x 11 | 5.544E-05 | 6.072 | 148.9 |
| 8.000 | 0.5 | USS | 15 x 15 | 5.555E-05 | 6.060 | 148.9 |
| 8.000 | 0.5 | USS | 21 x 21 | 5.578E-05 | 6.035 | 149.0 |
| 8.000 | 0.5 | USS | 41 x 41 | 5.707E-05 | 5.893 | 149.3 |

**Table S8**

| p | $\alpha$ | Sampling | resolution | $\det(\mathcal{I})$ | $\kappa$ | $\theta$ |
|---|---|---|---|---|---|---|
| 8.000 | 0.5 | URS | 7 x 7 | 8.518E-06 | 8.486 | 89.4 |
| 8.000 | 0.5 | URS | 11 x 11 | 8.549E-06 | 8.455 | 89.4 |
| 8.000 | 0.5 | URS | 15 x 15 | 8.594E-06 | 8.410 | 89.4 |
| 8.000 | 0.5 | URS | 21 x 21 | 8.688E-06 | 8.319 | 89.4 |
| 8.000 | 0.5 | URS | 41 x 41 | 9.240E-06 | 7.818 | 89.4 |
